# Transcriptomic Landscape and Immune Microenvironment Around Wound Bed Define Regenerative versus Non-regenerative Outcomes in Mouse Digit Amputation

**DOI:** 10.1101/2024.12.04.626910

**Authors:** Archana Prabahar, Connie S. Chamberlain, Ray Vanderby, William L. Murphy, William Dangelo, Kulkarni Mangesh, Bryan Brown, Barsanjit Mazumder, Stephen Badylak, Peng Jiang

## Abstract

In the mouse distal terminal phalanx (P3), it remains unclear why amputation at less than 33% of the digit results in regeneration, while amputation exceeding 67% leads to non-regeneration. Unraveling the molecular mechanisms underlying this disparity could provide crucial insights for regenerative medicine. In this study, we aim to investigate the tissues within the wound bed to understand the tissue microenvironment associated with regenerative versus non-regenerative outcomes. We employed a P3-specific amputation model in mice, integrated with time-series RNA-seq and a macrophage assay challenged with pro- and anti-inflammatory cytokines, to explore these mechanisms. Our findings revealed that non-regenerative digits exhibit a greater intense early transcriptional response in the wound bed compared to regenerative ones. Furthermore, early macrophage phenotypes differ distinctly between regenerative and non-regenerative outcomes. Regenerative digits also display unique co-expression modules related to Bone Morphogenetic Protein 2 (BMP2). The differentially expressed genes (DEGs) between regenerative and non-regenerative digits are enriched in targets of several transcription factors, such as HOXA11 and HOXD11 from the HOX gene family, showing a time-dependent pattern of enrichment. These transcription factors, known for their roles in bone regeneration, skeletal patterning, osteoblast activity, fracture healing, angiogenesis, and key signaling pathways, may act as master regulators of the regenerative gene signatures. Additionally, we developed a deep learning AI model capable of predicting post-amputation time and level from RNA-seq data, with potential applications in personalized treatment strategies and assessing the impact of interventions on regenerative outcomes.

## Introduction

Human limb regeneration is inherently restricted, with a limited native regenerative response. Numerous approaches, such as cell transplantation, immune modulation, gene manipulation, and tissue engineering, have been explored to improve these capabilities(Choi et al., 2014; Dolan et al., 2018; Gultian et al., 2022; Oliveira et al., 2021; Ren et al., 2019; Shieh and Cheng, 2015; Wang et al., 2023; Yu et al., 2012). Nonetheless, the success of these methods is frequently hindered by a lack of complete knowledge about the fundamental understanding of the underlying regenerative mechanisms. A comprehensive understanding of these mechanisms is crucial, as it could revolutionize the field by enabling a shift towards more effective regenerative responses, marking a paradigm shift in therapeutic approaches.

Studies using animal models, particularly mouse digit tip regeneration models, have illustrated the complexity of the regeneration process. For instance, mutation or absence of critical genes, such as Lmx1b(Castilla-Ibeas et al., 2023), can disrupt the regeneration process, leading to non-regenerative outcomes. These findings underscore the intricate nature of digit regeneration and indicate that disruption of essential components can eliminate the ability to regenerate. Despite these insights, it remains largely unclear how to induce regenerative capabilities in normally non-regenerative digits under typical conditions.

In the mouse distal terminal phalanx (P3), regenerative outcomes are mainly determined by the extent of amputation. Amputations removing less than 33% of the distal phalanx lead to complete regeneration, while those removing more than 67% generally result in non-regenerative outcomes(Borgens, 1982; Chamberlain et al., 2017; Fernando et al., 2011; Han et al., 2008; Neufeld and Zhao, 1995). This amputation level dependent regeneration phenomenon is analogous to digit tip regeneration in humans(Fleckman et al., 2013) and has an intrinsic advantage over studying other commonly used non-mammalian appendage regeneration models such as the salamander limb or zebrafish fin. However, the underlying mechanisms for these amputation level dependent regenerative capabilities remain mysterious.

In this study, we used a mouse distal terminal phalanx (P3) level specific amputation model combined with an *in vitro* macrophage polarization assay and RNA sequencing to compare regenerative and non-regenerative digits. Our study addressed the following questions: (a) What are the transcriptomic differences between regenerative and non-regenerative digits over time? (b) Do regenerative and non-regenerative digits exhibit distinct macrophage phenotypes following amputation? (c) Are there specific transcription factors that drive the transcriptomic differences observed between regenerative and non-regenerative digits post-amputation? (d) Does gene regulation vary depending on the regenerative or non-regenerative context?

Since the amputation level directly influences the likelihood of digit regeneration, then, predicting the amputation level can forecast the likelihood of regeneration. Additionally, the timing post-amputation is an important factor in measuring the healing process. By leveraging transcriptomic data, we have developed a deep learning (DL) model that can accurately predicts the level of amputation and the timing post-amputation. This model has the potential to improve our ability to tailor strategies and customize treatments for individual patients, ultimately enhancing clinical outcomes

## Results

### The regenerative capability of mouse distal terminal phalanx (P3) is amputation level dependent

Regeneration of the mouse digit is dependent on the initial level that is removed. If the less than 33% of the distal phalanx, including bone, ligaments, soft tissue, etc is amputated, the digit will regenerate. In contrast, removal of over 67% of the distal phalanx will result in a non-regenerating digit. **Figure 1A-D** illustrates the different amputation levels based on microCT scan: intact digits (**Figure 1A**), regenerative digits with amputation levels below 33% (**Figure 1B**; post-amputation day 14), intermediate regenerative digits with amputation levels between 33% and 67% (**Figure 1C**; post-amputation day 14), and non-regenerative digits with amputation levels exceeding 67% (**Figure 1D**; post-amputation day 14). Corresponding Hematoxylin and eosin (H&E) stained sections are shown in **Figure 1E-H**.

**Figure 1:**
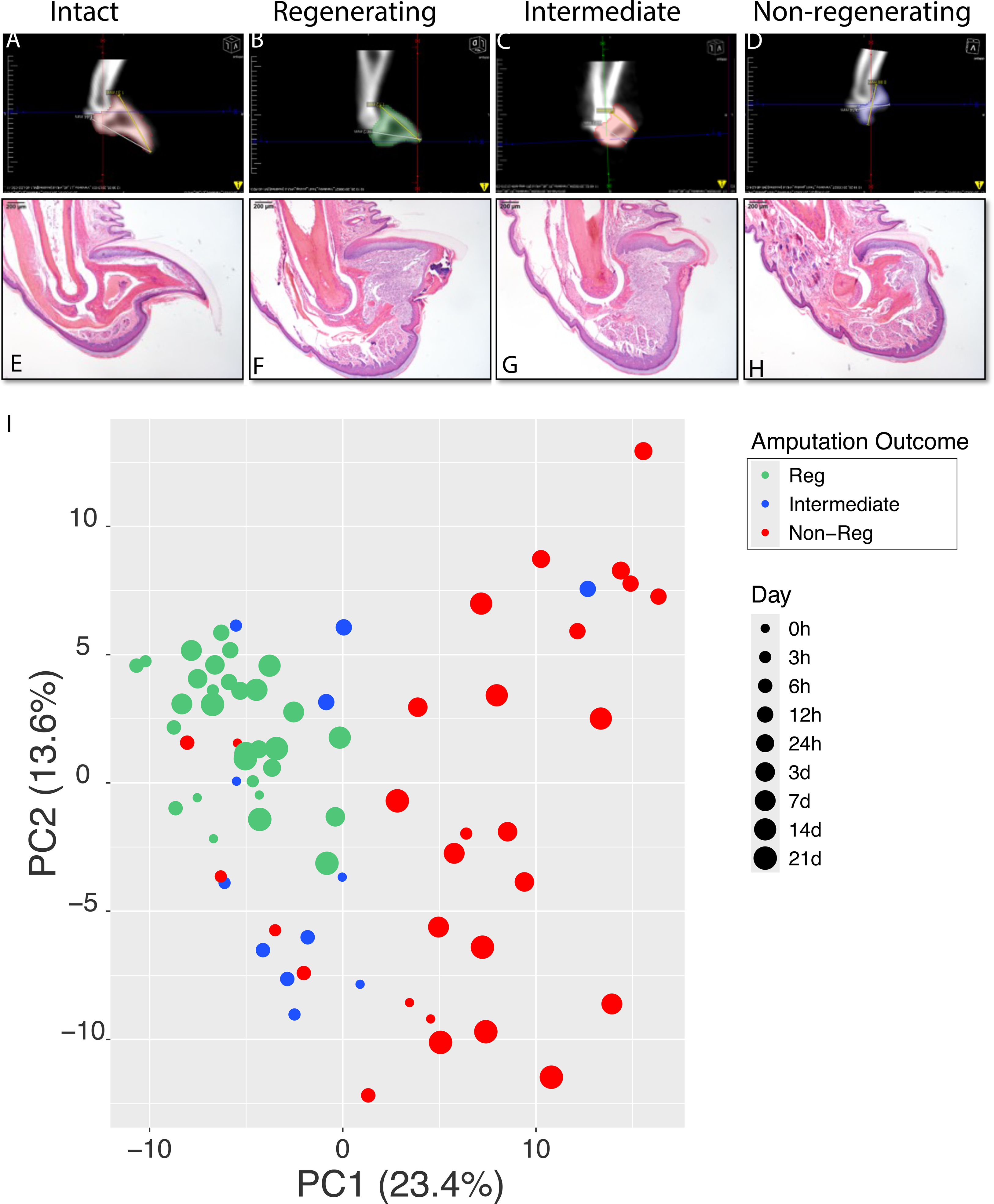
Amputation-Level-Dependent Regenerative Capacity of the Mouse Distal Terminal Phalanx. (A-D): MicroCT scans. (E-H): Hematoxylin and eosin (H&E) stained sections. (A) and (E): intact (no amputation); (B) and (F): regenerative digits (amputation level < 33%; post-amputation day 14); (C) and (G): intermediate amputation (amputation level 33–66%; post-amputation day 14); (D) and (H): non-regenerative digits (amputation level > 66%; post-amputation day 14); (I) Principal component analysis (PCA) of RNA-seq-derived gene expression in the wound bed post-amputation. Dot size represents the post-amputation time (days or hours).

We conducted RNA sequencing (RNA-seq) at various time points: immediately at the time of amputation (0 hours), and then at 3, 6, 12, 24 hours, and 3, 7, 14, and 21 days post-amputation. The level of amputation ranged from 10% in regenerative digits to 80% in non-regenerative digits. Our analysis focused on the transcriptomic similarities and differences across these wound bed tissues. As depicted in **Figure 1I**, the principal component analysis (PCA) clearly distinguishes regenerative digits (amputation level <33%) from non-regenerative digits (amputation level >67%), while the intermediate digits (amputation levels ranged from 34% to 67%) exhibit substantial overlap with both groups. This suggests that the intermediate digits may represent a transitional state between regenerative and non-regenerative stages.

### Non-regenerative digits exhibit more severe early transcriptional response to amputation compared to regenerative digits

To better understand the differences between regenerative and non-regenerative digits, we focused on the transcriptional activity post-amputation, as these patterns can provide critical insights into the molecular mechanisms driving regeneration. After amputation, we investigated whether transcriptional activity is consistently high or low, or exhibits fluctuations such as sudden bursts at specific time points. We noted a significant transcriptional burst in non-regenerative digits (amputation level >67%) within the first 12 hours post-amputation (**Figure 2A**), suggesting an immediate and strong early transcriptional response to injury. In contrast, regenerative digits (amputation level <33%) did not exhibit significant early transcriptional burst, showing only a marginal increase transcriptional activity compared to later time points (**Figure 2A**). This suggests that regenerative digits exhibit more stable gene expression patterns post-amputation, while non-regenerative digits show a stronger response to the injury.

**Figure 2:**
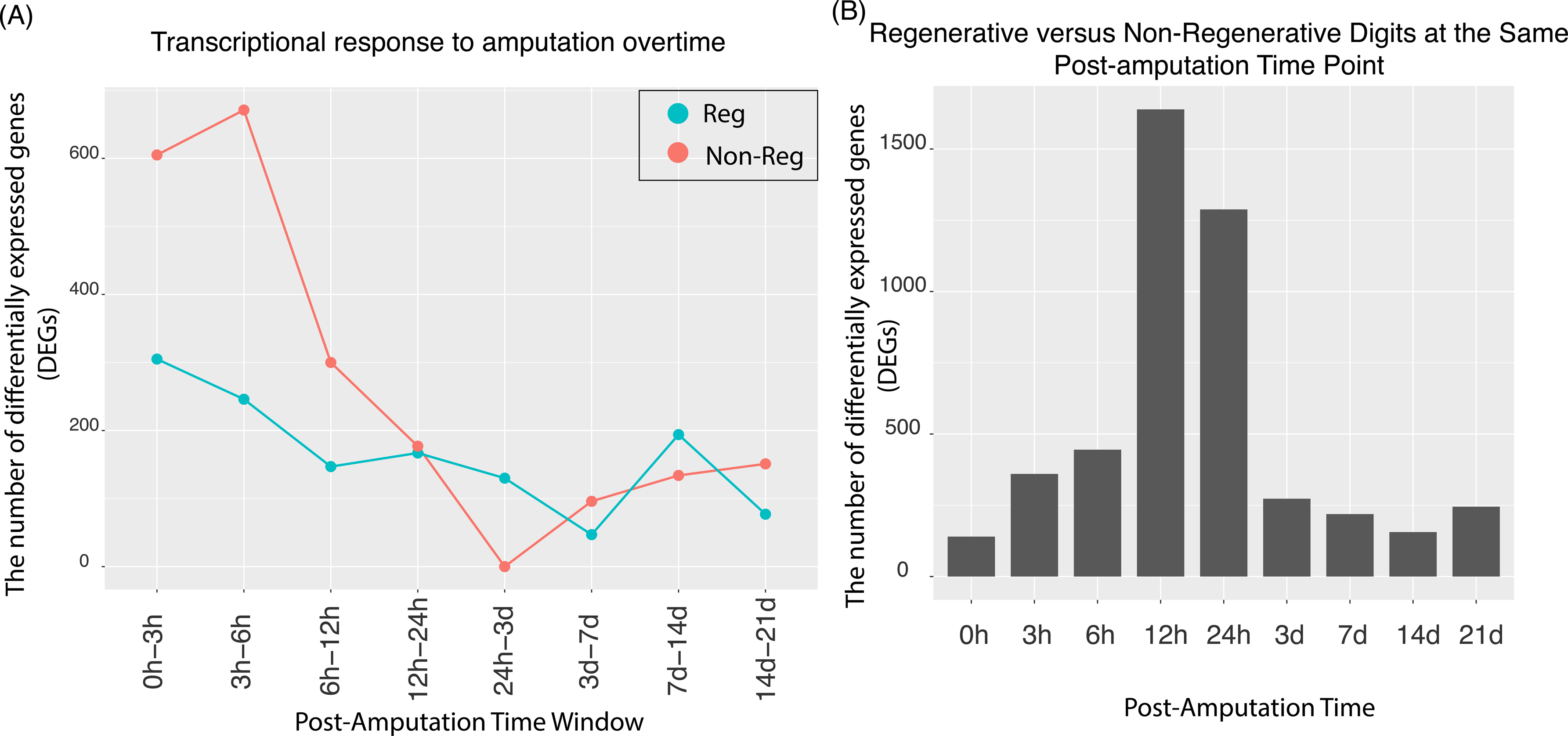
Differentially Expressed Genes (DEGs) in Regenerative vs. Non-Regenerative Digits. (A) DEGs identified at various time intervals following amputation; (B) DEGs comparing regenerative and non-regenerative digits at the same post-amputation time point.

We further analyzed the number of genes differentially expressed between regenerative and non-regenerative digits at the same post-amputation time point. As shown in **Figure 2B**, there was minimal transcriptomic difference between regenerative and non-regenerative digits at the time of amputation (0 hour), indicating that inherent positional differences between regenerative and non-regenerative digits are minimal. This suggests that the differences in healing outcomes (regenerative versus non-regenerative) are primarily driven and initiated by changes that occur post-amputation (e.g., cell migration, proliferation and differentiation) rather than by pre-existing positional differences. From 12 to 24 hours post-amputation, both regenerative and non-regenerative digits exhibited a similar number of differentially expressed genes (**Figure 2A**). However, these time points marked the period with the most significant transcriptomic differences between the regenerative and non-regenerative digits (**Figure 2B**). This suggests that regenerative and non-regenerative digits follow distinct transcriptomic activity patterns post-amputation, and the major differences peaks at early time points (e.g., ∼ 12-24 hours post-amputation).

The significant early transcriptional burst observed in non-regenerative digits highlights a more severe initial response to amputation compared to regenerative digits. The highest number of differentially expressed genes between regenerative and non-regenerative digits occurs at early post-amputation time points. These early transcriptional differences underscore the critical role of initial molecular responses in determining the divergent healing outcomes between regenerative and non-regenerative digits.

### Macrophage polarization patterns in early time points distinguish the regenerative and non-regenerative outcomes

In mammals, macrophages are known to play a major role in tissue regeneration(Lavin and Merad, 2013; Simkin et al., 2017; Yu et al., 2022). The general notion is that tissue regeneration follows a biphasic inflammatory pattern where the early phase is dominated by pro-inflammatory signals and macrophages exhibit a classically activated, or M1, phenotype. These M1 macrophages release cytokines and other factors that are crucial for clearing pathogens and debris, thus setting the stage for healing. As the regenerative process progresses, there is a shift towards an anti-inflammatory environment, where alternatively activated, or M2, macrophages predominate. These M2 macrophages facilitate tissue repair and regeneration by secreting anti-inflammatory cytokines and growth factors, supporting angiogenesis, and remodeling the extracellular matrix. Thus, the transition from a pro-inflammatory to an anti-inflammatory phase is critical for successful tissue regeneration(Landen et al., 2016). Previous studies have shown that depletion of macrophages will completely repress the mouse digit tip regeneration(Simkin et al., 2017). However, the role of macrophage polarization to modulate the digit regeneration outcomes remains uncertain—for instance, it’s unclear whether regenerative digits undergo a different macrophage polarization trajectory compared to non-regenerative digits. Assessing the role of macrophage polarization in digit regeneration is technically challenging. This is due to the fact that most of the canonical M1 and M2 markers are context-dependent(Strizova et al., 2023). To address potential biases in defining M1-like or M2-like macrophages based solely on a few canonical markers, we adopted an *in vitro* approach to refine these classifications. We exposed bone marrow-derived macrophages to pro-inflammatory cytokines (IFN-γ and LPS) and anti-inflammatory cytokines (IL-4), and compared their responses to control (M0) macrophages. Through RNA-seq, we systematically identified gene signatures characteristic of M1-like and M2-like states. As illustrated in **Figure 3A**, we identified 2,728 M1 and 1,355 M2 macrophage gene markers, up-regulated in macrophages stimulated by IFN-γ/LPS and IL-4, respectively. This comprehensive list of M1 and M2 markers is used to assess macrophage polarization during the digit regeneration process (comparing regenerative and non-regenerative digits). This approach provides an unbiased and robust comparison, unlike the use of a few canonical M1 and M2 markers. We compared these markers with genes up-regulated in regenerative and non-regenerative digits post-amputation over time, relative to the time of amputation, to discern the specific macrophage phenotypes (M1 or M2) active during different phases of digit regeneration. Notably, non-regenerative digits (amputation level >67%) exhibited a sharp increase in M1 (pro-inflammatory) macrophage activation within the first 3 days (**Figure 3B**), followed by a rapid decline in M1 signals. In contrast, regenerative digits (amputation level < 33%) showed significantly lower levels of M1 macrophage activity in the first 3 days but in a very similar level at later time points. Interestingly, the M2 (anti-inflammatory) macrophage signals mirrored the M1 signals in both regenerative and non-regenerative digits (**Figure 3C**). We then calculated the M1/M2 ratio—the ratio of up-regulated genes post-amputation overlaid with M1 and M2 markers (**Figure 3D**). Our findings reveal a distinct difference between regenerative and non-regenerative digits at early post-amputation time points from 6 hours to day 3, where regenerative digits display a lower M1/M2 ratio, suggesting reduced inflammation during this critical early window, which may facilitate more effective regeneration.

**Figure 3:**
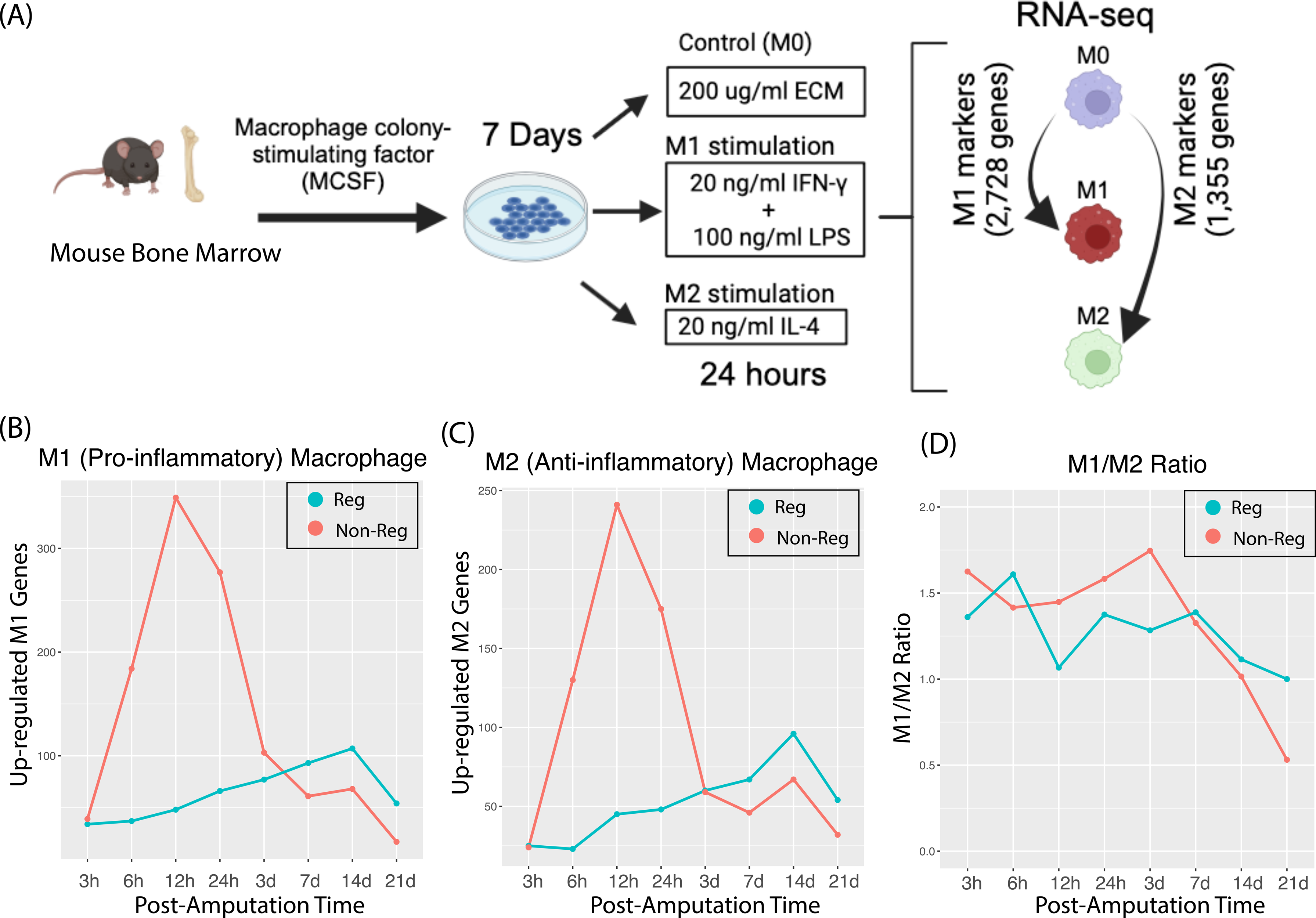
Macrophage Polarization Patterns Post-Amputation. (A) Workflow for identifying gene markers associated with pro-inflammatory (M1) and anti-inflammatory/pro-regenerative (M2) macrophages; (B) Number of upregulated genes overlapping with M1 (pro-inflammatory) markers in regenerative and non-regenerative digits relative to intact controls; (C) Number of upregulated genes overlapping with M2 (anti-inflammatory) markers in regenerative and non-regenerative digits relative to intact controls; (D) M1/M2 ratio (number of upregulated genes overlapping with M1 markers divided by those overlapping with M2 markers) in regenerative and non-regenerative digits compared to intact controls.

### Identifying biological process distinguish regenerative and non-regenerative digits

For each time point, we identified genes that were either up-regulated or down-regulated in regenerative digits compared to non-regenerative digits. We then performed a Gene Ontology (GO) enrichment analysis, and identified a broad spectrum of biological processes enriched in these differentially expressed genes (see **Supplementary Table S1 and S2**: enriched GO terms for up- and down-regulated genes, respectively). We further prioritized these enriched biological processes based on their enrichment (adjust P-values < 0.05) consistency across multiple time points.

Notably, for genes up-regulated in regenerative digits, five GO terms were consistently enriched (adjust P-values < 0.05) across all time points: BMP signaling pathway **(Figure 4A)**, Cellular response to BMP stimulus **(Figure 4B)**, Ossification **(Figure 4C)**, Osteoblast differentiation **(Figure 4D)**, and Response to BMP **(Figure 4E)**. The BMP signaling pathway, critical for digit regeneration, enhances osteoblast differentiation and bone formation, and supports chondrogenesis, angiogenesis, and interactions with vital pathways such as Wnt and FGF. The consistent enrichment of the BMP associated pathway regenerative digits underscores their importance in regeneration. Ossification and osteoblast differentiation, crucial for bone formation, are consistently enriched in the up-regulated genes of regenerative digits compared to non-regenerative ones in all time points. This suggests that digits with less than 33% amputation are actively involved in bone tissue regeneration, demonstrating a strong regenerative capability.

**Figure 4:**
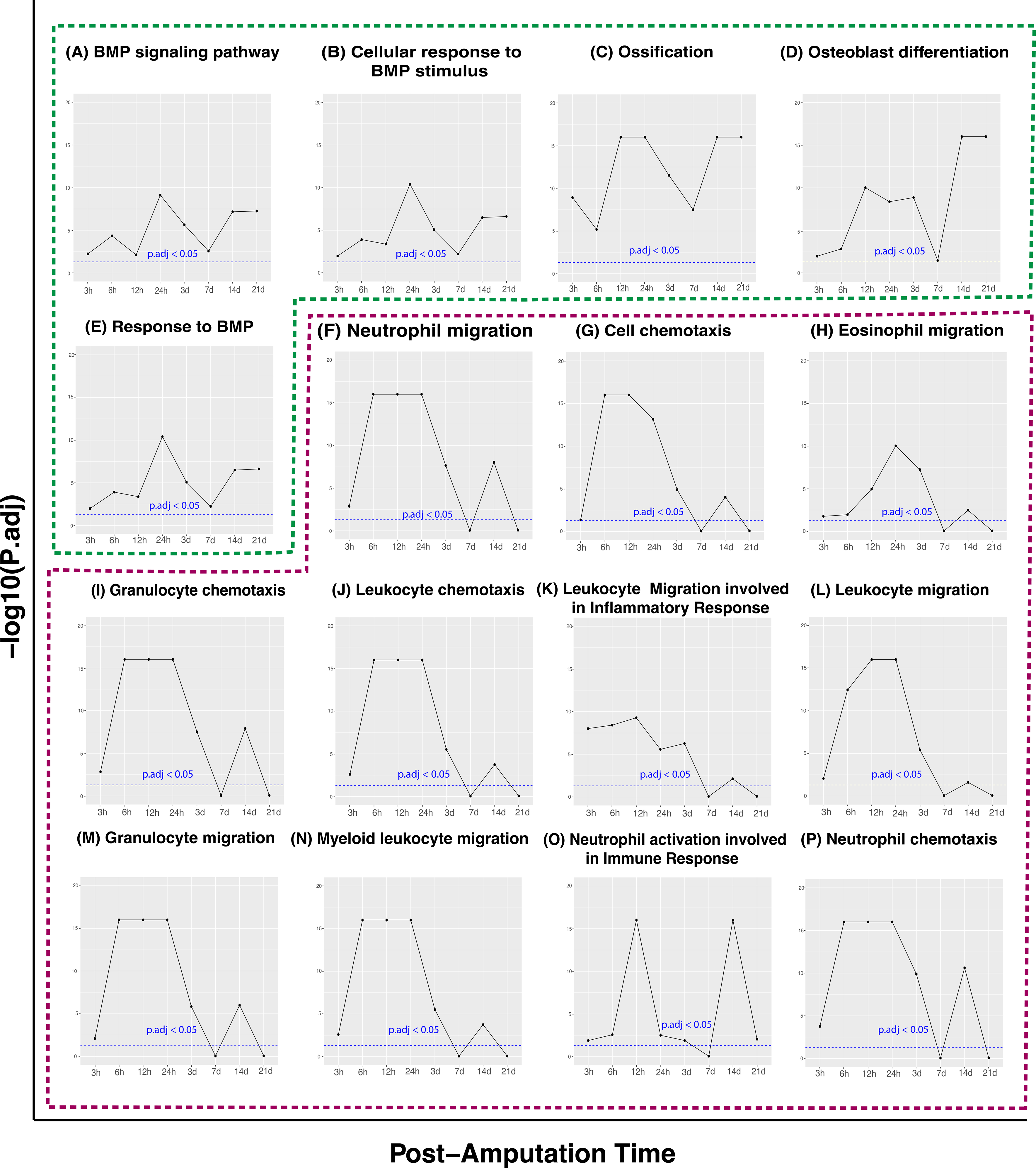
Consistently enriched GO terms across post-amputation time points comparing regenerative versus non-regenerative digits. Panels (A–E) highlight the top five GO terms that remain consistently enriched (adjusted p-values < 0.05) across all time points. Panels (F–P) display the top-ranked consistently enriched GO terms for downregulated genes, with rankings based on the number of post-amputation time points showing significant enrichment (adjusted p-values < 0.05).

In contrast, the most consistently enriched GO terms for genes down-regulated in regenerative digits across time points were related to immune responses **(Figure 4F-P)**, such as Leukocyte migration involved in inflammatory response, Neutrophil chemotaxis, Neutrophil migration, Granulocyte chemotaxis, Myeloid leukocyte migration, Leukocyte chemotaxis, Granulocyte migration, Leukocyte migration, Eosinophil migration, Cell chemotaxis, and Neutrophil activation involved in immune response. This further suggest that non-regenerative digits exhibit heightened inflammatory responses.

### Regenerative and non-regenerative digits exhibit distinct Bmp2 associated co-expression modules

Bone Morphogenetic Protein 2 (BMP2), a key member of the Transforming Growth Factor-Beta (TGF-β) superfamily, plays a pivotal role in both developmental biology and osteogenesis. BMP-2 is extensively involved in numerous developmental processes including digit formation and osteogenesis(Halloran et al., 2020; Wang et al., 2023; Yu et al., 2012). Multiple studies have shown that delivery BMP2 can promote bone regeneration(Gultian et al., 2022; Jeon et al., 2022). We investigated whether regenerative digits tend to have higher expression of Bmp2 compared to non-regenerative digits. Surprisingly, we did not observe a statistically significant increase in Bmp2 expression in regenerative digits at any of the post-amputation time points. There was only marginally higher expression in regenerative digits from day 3 to day 21, with differential expression p-values ranging from 0.12 (day 7) to 1 (24 hours) (**Supplementary Figure S1**). This indicates that Bmp2 is not a major factor distinguishing regenerative from non-regenerative digits. Given Bmp2’s role in bone remodeling and homeostasis(Halloran et al., 2020), we further investigated whether Bmp2-associated co-expression modules (groups of genes with similar expression patterns, which tend to be functionally related) differ between regenerative and non-regenerative contexts. We identified 294 genes co-expressed with Bmp2 in regenerative digits and 626 in non-regenerative digits (Spearman’s Rho > 0.5). Interestingly, only 42 genes were common between the two sets (**Figure 5**). Gene Ontology (GO) enrichment analysis of these genes revealed distinct top enriched GO terms in regenerative versus non-regenerative digits (**Figure 5**). This indicates that the BMP2-associated co-expression modules are markedly different in regenerative compared to non-regenerative digits, suggesting a potential Bmp2 associated unique regulatory mechanisms depending on the regenerative capacity of the digits.

**Figure 5:**
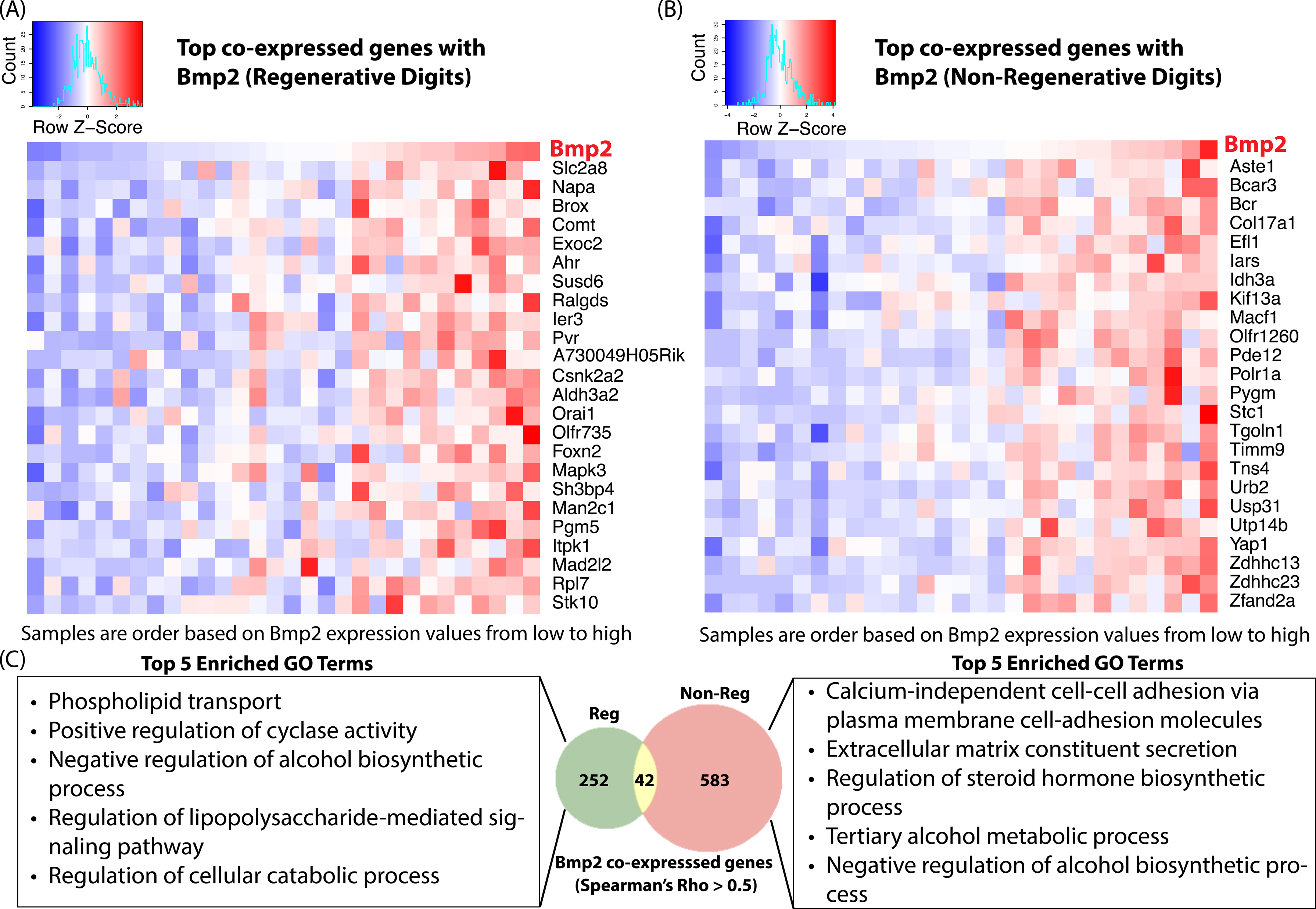
Context-dependent co-expression of Bmp2 gene. (A) Top co-expressed genes with Bmp2 in regenerative digits; (B) Top co-expressed genes with Bmp2 in non-regenerative digits; (C) Overlapping of top co-expressed genes with Bmp2 in regenerative versus non-regenerative digits, and enriched GO terms.

### Prioritizing key regulators responsible for transcriptomic variations between regenerative and non-regenerative digits

In a comparison of regenerative and non-regenerative digits, there is a differential expression of an average of 578 genes at the same post-amputation time points, with variations ranging from 156 DEGs at day 14 to 1639 DEGs at 12 hours post-injury. Overall, 2984 genes show differential expression at any given post-amputation time point. To pinpoint potential master regulators driving these differences, we conducted a transcription factor enrichment analysis (see Methods).

In regenerative digits, compared to non-regenerative ones, 52 transcription factors exhibited enrichment in up-regulated genes, as shown in **Figure 6A**. These transcription factors demonstrate patterns of enrichment that vary over time. Notably, at 12 hours post-amputation, HOXA11 and HOXD11 are enriched; these factors are pivotal for chondrocyte differentiation and bone development. Additionally, TBX4 and EN1, which are essential for re-establishing limb identity and growth patterns, as well as re-establishing apical ectodermal ridge (AER)-like signaling centers crucial for limb outgrowth and patterning, respectively, do not show enrichment until day 21.

**Figure 6:**
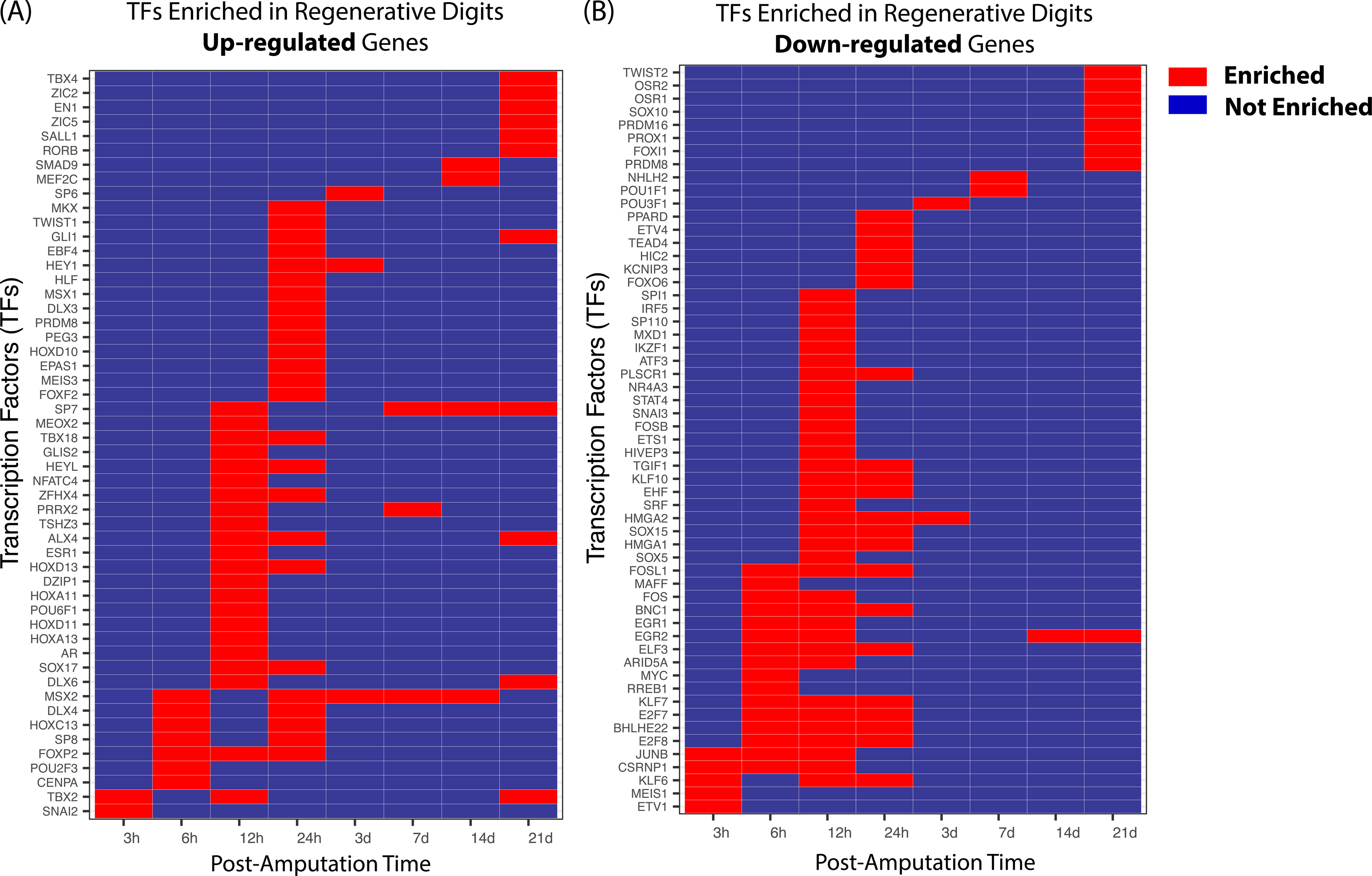
Transcription factor (TF) enrichment analysis identifying potential TFs associated with differentially expressed genes (DEGs) between regenerative and non-regenerative digits. (A) TFs whose target genes are enriched among upregulated DEGs in regenerative digits compared to non-regenerative digits. (B) TFs whose target genes are enriched among downregulated DEGs in regenerative digits compared to non-regenerative digits.

In the analysis of down-regulated genes in regenerative digits compared to non-regenerative ones, 57 transcription factors were found to be enriched (**Figure 6B**). Among these, JUNB, a member of the AP-1 transcription factor family, was notably enriched at early post-amputation time points (within the first 24 hours). JUNB’s target genes are down-regulated in regenerative digits during early post-amputation time points (within the first 24 hours), but this enrichment is not observed at later stages. JUNB binds the COL1A2 enhancer, driving the transcription of type I collagen, a major component of extracellular matrix deposition(Ponticos et al., 2015). Studies have shown that reducing JUNB expression decreases activation of the COL1A2 enhancer and subsequent type I collagen production(Ponticos et al., 2015), underscoring JUNB’s role in promoting collagen overproduction, a process linked to pro-fibrotic activity. While type I collagen is crucial for structural integrity, particularly in bones where it provides a framework for mineral deposition and strength, excessive deposition is a hallmark of fibrosis. These findings suggest that in the early post-amputation period, overexpression of JUNB may negatively impact digit regeneration by promoting a fibrotic response, hindering proper tissue remodeling and repair.

### Deep learning model to impute amputation level and post-amputation time

The amputation level directly influences the likelihood of digit regeneration, with favorable outcomes typically occurring when the amputation level is under 33%, while non-regenerative outcomes are more common beyond 67%. Thus, predicting the amputation level can forecast the likelihood of regeneration. Post-amputation time serves as another critical indicator of healing progress. By leveraging transcriptomic data to predict these parameters, we can anticipate whether injured digits are likely to regenerate or follow a non-regenerative path. Predicting both post-amputation time and amputation level together enhances our understanding of digit injury recovery and informs potential interventions.

However, jointly predicting post-amputation time and amputation level presents technical challenges due to their interdependence. For instance, the transcriptomic profile at a specific post-amputation time may differ between regenerative and non-regenerative digits. To address this, we initially developed independent random forest (RF) models to identify genes associated with either post-amputation time or amputation levels (**Figure 7A**). We identified 149 genes that account for over 95% of the feature importance contribution of either of the random forest models. Notably, these genes were enriched in pathways related to extracellular matrix degradation, collagen degradation, and tissue organization, as well as in Gene Ontology terms such as morphogenesis of anatomical structures, tissue morphogenesis, and epithelium development (**Figure 7A**). Subsequently, these 149 genes were used as inputs to construct a deep learning model with two hidden layers, aiming to predict both post-amputation time and amputation levels simultaneously (**Figure 7B-C**). The deep learning model demonstrated high accuracy in predicting these parameters, achieving Spearman’s Rank correlation coefficients (Rho) of 0.87 and 0.92 for predicted versus observed amputation level and post-amputation time, respectively.

**Figure 7:**
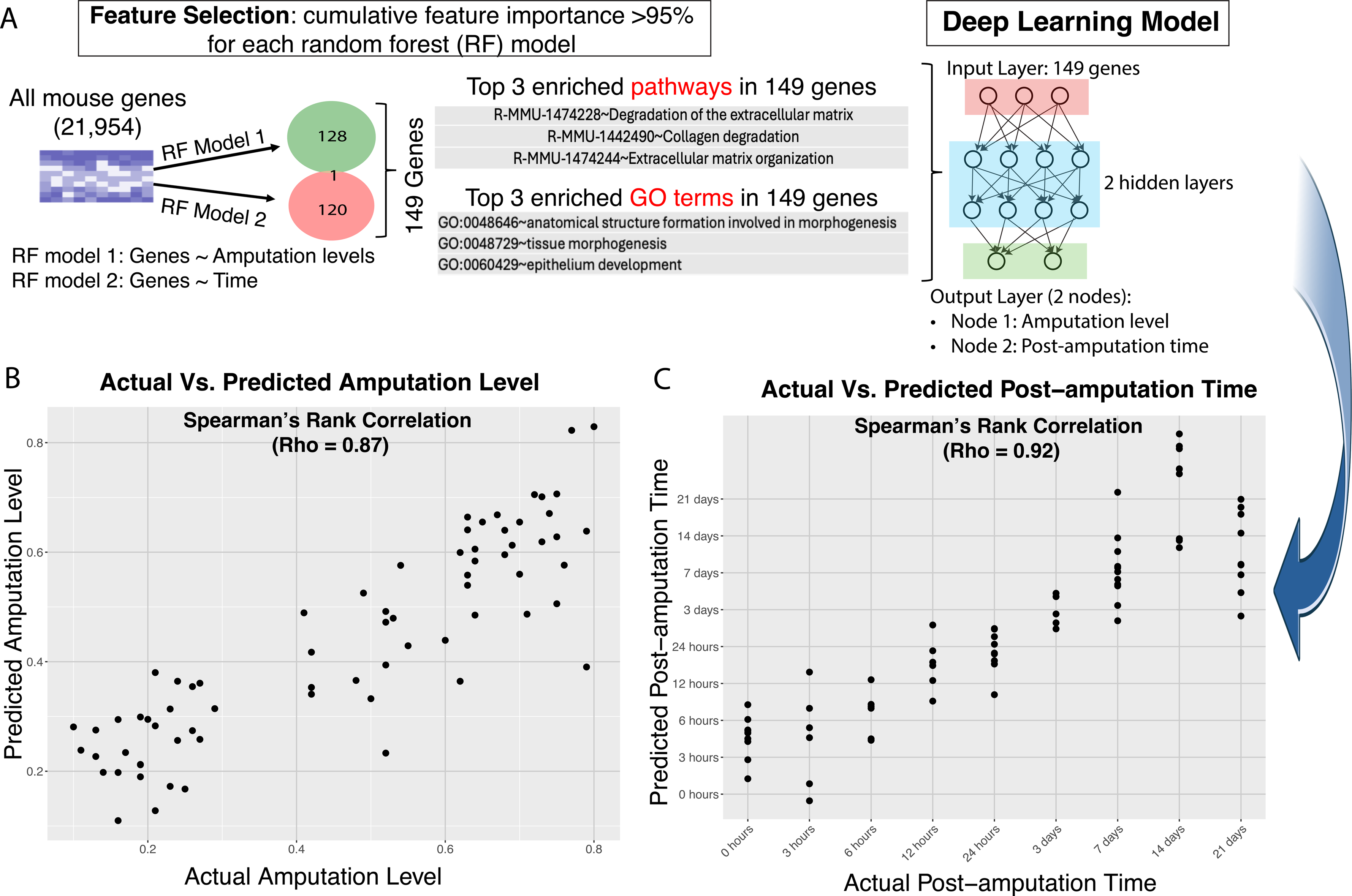
Deep learning (DL) model to impute amputation level and post-amputation time based on wound bed gene expression patterns. (A) feature selection via a random forest model; (B) Five-fold cross-validation for DL model to predict amputation level; (C) Five-fold cross-validation for DL model to predict post-amputation time.

## Discussion

The mechanisms by which amputations below 33% of the mouse distal terminal phalanx (P3) facilitate regeneration, whereas those above 67% fail to do so, are not fully understood but are essential for progressing regenerative medicine. Our study underscores the significance of initial post-amputation responses in shaping these outcomes. Specifically, (a) non-regenerative digits exhibit a substantially more intense early transcriptional response than regenerative ones, underscoring the importance of early molecular activities; (b) The initial polarization patterns of macrophages predict the potential for digit regeneration. These findings indicate that the capacity for regeneration may be determined very early post-amputation, emphasizing the early period as crucial. This insight opens a potential window for therapeutic intervention aimed at targeting these initial phases to enhance regenerative outcomes.

Bmp2, a critical component of the TGF-β superfamily, plays important role in digit formation and osteogenesis(Halloran et al., 2020; Wang et al., 2023; Yu et al., 2012). Although multiple studies suggest that Bmp2 administration can enhance bone regeneration(Gultian et al., 2022; Jeon et al., 2022), our findings did not reveal significant differences in BMP-2 expression between regenerative and non-regenerative digits post-amputation. This suggests that while Bmp2 can potentially enhance digit regeneration under specific conditions, it is not alone capable of converting non-regenerative responses into regenerative ones. Interestingly, genes that are significantly up-regulated in regenerative digits compared to non-regenerative ones are enriched in BMP-associated GO terms, such as Response to BMP (see **Figure 4E**). Further investigation also reveals that regenerative and non-regenerative digits display distinct Bmp2-associated co-expression modules. This indicates that while Bmp2 expression levels do not differ significantly between regenerative and non-regenerative digits, the regulatory context or gene regulatory networks related to Bmp2 differ substantially between these two post-amputation outcomes. This suggests a complex role for Bmp2 in influencing digit regeneration that goes beyond mere expression levels.

The capacity for digit regeneration is influenced by the level of amputation: less than 34% typically results in regeneration, more than 67% in non-regeneration, and amputation between these percentages shows variable or partial regenerative abilities. At the transcriptomic level, we investigated whether intermediate amputations exhibit a unique pattern or overlap with regenerative or non-regenerative digits. Our findings indicate that while regenerative and non-regenerative digits display distinct patterns, intermediate amputations share similarities with both, suggesting a gradual rather than binary transition in the transcriptomic landscape related to amputation level. This implies that the regenerative potential of a digit could be quantified in terms akin to “amputation level” values. Additionally, the post-amputation time serves as an indicator of the transcriptomic state of the wound. Thus, amputation level and post-amputation time are critical metrics for assessing potential outcomes (regeneration or non-regeneration) and the status of wound healing. We have developed a deep learning model that predicts the amputation level and the elapsed time since amputation using transcriptome data. This model offers potential benefits for guiding personalized treatment strategies and enhancing outcomes in regenerative medicine by enabling precise interventions based on the specific conditions of a digit injury. By predicting the injury conditions, such as the level of amputation or the time since injury, appropriate therapeutic measures can be tailored more accurately. Furthermore, this model also holds the potential to assess the effectiveness of interventions, such as whether they accelerate the regeneration process or shift the transcriptomic profile towards a regenerative outcome from a non-regenerative state.

## Methods

### Mouse distal terminal phalanx (P3) level-specific amputations comparative regenerative model

All experiments performed were approved by the University of Wisconsin-Madison Institutional Animal Care and Use Committee. A total of 53 male C57Bl/6 mice (9-10 week old) were included in the study. Animals were subjected to hindlimb bilateral distal phalanx (P3) amputation of digit 3. For every amputation, mice were anesthetized using isoflurane, the hindlimb claw was extended, and the distal phalanx and footpad was sharply dissected. A regenerating distal phalanx was created by amputating ≤33% of the ipsilateral P3 whereas a non-regenerating distal phalanx was generated by amputating > 67% of the contralateral P3. An “intermediate” regenerating digit was created by amputating 34-66% of P3. Skin wounds were allowed to heal without suturing. To confirm the length of distal phalanx was accurately removed, mouse digits were subjected to micro-computed tomography (microCT) prior to surgery and immediately after amputation. Any digit falling outside the amputation guidelines was removed from the study. Based on this criteria, 14 animals were removed from the study resulting in a final total of 39 mice (3 mice/day). At 0h, 3h, 6h, 12h, 24h, 3d, 7d, 14d, and 21d the regenerating and non-regenerating distal phalanges were collected.

Intermediate digits were collected at 0h, 24h, 14d, and 21d post-amputation. Another group of mice did not undergo amputation and digits served as the intact controls. At the time of P3 collection, digits were immediately immersed in RNAlater for 24h at 4C. The samples were then removed from RNAlater and stored at -80C until RNA-seq. Total RNAs were isolated from tissues using trizol (ThermoFisher #15596018) and chloroform phase separations followed by the RNeasy mini protocol (Qiagen #74106) with optional on-column DNase digestion (Qiagen #79254). One hundred nanograms of total RNA was used to prepare sequencing libraries using the LM-Seq (Ligation Mediated Sequencing) protocol(Hou et al., 2015). RNA was selected using the NEB Next Poly A+ Isolation Kit (NEB #E7490S/L). Poly A+ fractions were eluted, primed, and fragmented for 7 minutes at 85°C. First strand cDNA synthesis was performed using SmartScribe Reverse Transcriptase (Takara Bio USA #639538) and RNA was removed. cDNA fragments were purified with AMpure XP beads (Beckman Coulter #A63881). The 5’ adapter was ligated and 18 cycles of amplification were performed. These final indexed cDNA libraries were quantified, normalized, multiplexed, and run as single-end reads for 65bp on the HiSeq 2500 (Illumina, San Diego, CA).

#### MicroCT Analysis

Mice hindlimb paws were longitudinally imaged using microCT to assess digit regeneration. MicroCT provides the necessary resolution and contrast to measure digit length and volume used in the analysis. Imaging was accomplished via Siemens Inveon microCT scanner, and analysis was conducted using Inveon Research Workplace General and 3D Visualization software (Siemens Medical Solutions USA, Inc., Knoxville, TN). CT scans were acquired with the following parameters: 80 kVp, exposure time, 900 µA current, 220 rotation steps with 441 projections, ∼16.5 minute scan time, bin by 2, 50 micron focal spot size, and medium magnification that yielded an overall reconstructed isotropic voxel size of 46.6 um^3^. Raw data were reconstructed using filtered back-projection and no down-sampling via integrated high-speed COBRA reconstruction software (Exxim Computing Corporation, Pleasanton, CA). Hounsfield units (HU), a scalar linear attenuation coefficient, was applied to each reconstruction to permit inter-subject comparisons. Three-dimensional images were segmented using a pixel intensity range between 300 HU (minimum) and 3168 HU (maximum) to represent bone density. Once the region of interest (ROI) was defined, the P3 volume was calculated. Sagittal digit length was also obtained by measuring twice from the distal tip to proximal edge of the P3 bone. Two blinded, independent researchers conducted the digit measurements, and their results were averaged.

### Murine Bone Marrow-Derived Macrophage Isolation and Treatment

Bone marrow-derived macrophages (BMDMs) were isolated and characterized from 6– 8-week-old female C57BL/6 mice, following the protocol described by (Sicari et al., 2014). Mice were euthanized using CO₂ and cervical dislocation, and tibias and femurs were aseptically harvested. Bone marrow cells were flushed from the bones using a 30-gauge needle and macrophage complete medium (high glucose DMEM supplemented with 10% fetal bovine serum [FBS], 10% L929 conditioned medium, 0.1% β-mercaptoethanol, 100 U/ml penicillin, 100 U/ml streptomycin, 10 mM non-essential amino acids, and 10 mM HEPES buffer). The cells were then washed, plated at a density of 10⁶ cells/ml, and cultured for 7 days in complete medium, with medium changes every 48 hours. On day 7, naive macrophages (M0) were treated with basal medium (high glucose DMEM supplemented with 10% FBS, 100 U/ml penicillin, and 100 U/ml streptomycin). For polarization, M0 macrophages were treated in basal medium as follows:

- M1 phenotype: 20 ng/ml IFN-γ + 100 ng/ml LPS
- M2 phenotype: 20 ng/ml IL-4

After 24 hours of treatment, cells were collected for RNA isolation and performed RNA-seq.

### RNA-seq data analysis

RNA-seq Reads were mapped to the mouse genes (Ensembl v75) using Bowtie (v0.12.8) (Langmead et al., 2009) allowing up to 2-mismatches and a maximum of 200 multiple hits. The gene expected read counts and TPMs were estimated by RSEM (v1.2.3) (Li and Dewey, 2011). TPMs were median-by-ratio normalized(Leng et al., 2013), and replicates were merged via calculating average normalized TPMs.

### Differentially expressed genes (DEGs)

The EBSeq package(Leng et al., 2013) was used to assess the probability of gene expression (mRNAs) being differentially expressed between any two given conditions. We required that DEGs should have FDR<5% via EBSeq and >2 fold-change of “normalized read counts+1”.

### Gene Ontology (GO) enrichment analysis

Gene ontology (GO) enrichment analysis was performed using the R package (“allez”) (Newton et al., 2007). The enrichment p-values were further adjusted by Benjamini– Hochberg (BH) multiple test correction.

### Principal Component Analysis (PCA)

To determine whether the intermediate amputated digits (amputation levels are from 34% to 67%) are more transcripomic similar to regenerative (amputation level <33%) or non-regenerative digits (amputation level >67%), we conducted a Principal Component Analysis (PCA). Considering that various factors can influence the transcriptome, we first identified differentially expressed genes between regenerative digits and non-regenerative digits at each time point. We then selected genes that were differentially expressed (FDR<5% and >2-fold changes of “normalized expected counts +1”) between these two groups at any time point. This list of genes was used to perform the PCA across all three categories: non-regenerative digits, regenerative digits, and intermediate digits. Normalized TPM values were used for the PCA calculation.

### Transcription factor enrichment analysis

To pinpoint the transcription factors potentially responsible for the differential gene expression between regenerative and non-regenerative digits, we conducted a transcription factor enrichment analysis. For each post-amputation time point, up-regulated and down-regulated genes (regenerative vs non-regenerative digits) were identified as those with an FDR < 5% using EBSeq(Leng et al., 2013) and exhibiting a fold-change >2 in “normalized read counts +1. We then analyzed these up-regulated and down-regulated gene sets separately to identify transcription factors whose target genes were significantly enriched within each list. ChEA3(Keenan et al., 2019) tool was used to identify potential TFs whose targets are enriched in the up or down regulated genes. In brief, ChEA3 leverages a comprehensive database of transcription factor-target interactions derived from ChIP-seq, ChIP-chip, and other sources. ChEA3 calculates the overlap between the input gene list and known transcription factor targets, assessing significance with Fisher’s Exact test. We performed transcription factor enrichment analysis for up-regulated and down-regulated genes separately. Additionally, we required that enriched transcription factors (Fisher’s Exact Test p-value <0.05) based on the up-regulated or down-regulated gene lists should themselves exhibit corresponding up- or down-regulation (FDR < 5% and >2-fold changes), aligning with the expectation that transcription factors act as activators of gene expression. This requirement ensures that the identified transcription factors are not only enriched in their targets but are also likely playing an active regulatory role in driving the observed differential gene expression patterns.

### Deep learning model to impute amputation level and post-amputation time from transcriptome data

We developed a deep learning (DL) model to predict both the amputation level and post-amputation time, leveraging its significant advantage of being able to integrate multiple outputs within a single framework. Unlike traditional machine learning approaches, such as support vector machine (SVM)(Awad et al., 2015), which require separate models for each output, deep learning model(Goodfellow et al., 2016) can simultaneously predict multiple related parameters. This integration is particularly useful in our context, where amputation level and post-amputation time are interrelated but independent variables.

#### Feature selection

To select the most relevant genes as features for our deep learning model, we first employed a random forest model(Liaw and Wiener, 2002)—a machine learning technique that uses an ensemble of decision trees to enhance prediction accuracy and reduce overfitting. In this case, because we had two distinct outputs, the random forest model built two separate models: one for predicting amputation levels and another for post-amputation time. This process identified 128 genes associated with amputation levels and 120 genes associated to post-amputation time, based on their importance ranking. We selected genes whose cumulative contribution exceeded 50% of the total feature importance. Interestingly, there was minimal overlap between the genes associated with the two outputs, with only one gene common to both sets. This resulted in a final list of approximately 149 genes associated with either amputation levels, post-amputation time, or both, which were then used as input features for the deep learning model.

#### Deep learning model architecture

The deep learning model was implemented using TensorFlow(Abadi et al., 2019) and Keras(Gulli and Pal, 2017) packages. The model architecture consisted of an input layer consisting 149 nodes corresponding to the number of genes used as features, two hidden layers, and an output layer with two nodes (amputation level and post-amputation time). To optimize the model’s parameters, we implemented a systematic grid search approach. This involved testing various numbers of units in the first and second hidden layers to find the most effective configuration. Each configuration used the rectified linear unit (ReLU)(Agarap, 2018) for the hidden layers and a linear activation function for the output layer. The optimization goal was to maximize the product of Spearman’s Rho correlations for observed versus predicted values of both amputation level and post-amputation time. We employed a 5-fold cross-validation strategy to train and evaluate the model, ensuring robust performance and generalizability.

## Availability of data and materials

We have submitted mouse digit amputation wound bed RNA-seq data (<33% amputation level) to GEO with accession number: GSE130438. In this study, we submitted mouse digit amputation wound bed RNA-seq data (>34% amputation level) and macrophage in vitro (M0, M1 and M2 phenotypes) RNA-seq data to GEO with accession number: GSE279540.

## Supporting information

List_of_Supplementary_Data

## Acknowledgments

We thank the support from the Center for Gene Regulation in Health and Disease (GRHD) at Cleveland State University and Prof. Anton Komar. We thank the support from the Center for Gene Regulation in Health and Disease (GRHD) at Cleveland State University and Prof. Anton Komar. Thanks to the Wisconsin Institutes for Medical Research (WIMR) small animal vivarium for support. This work was also supported by DARPA AWD00001593 (PJ).

## Notes

### Competing Interest Statement

The authors have declared no competing interest.

