## Supplementary material for "Transcriptomic Landscape and Immune Microenvironment Around Wound Bed Define Regenerative versus Non-regenerative Outcomes in Mouse Digit Amputation": List_of_Supplementary_Data

### Supplementary Tables

**Supplementary Table S1:** Enriched GO terms for genes up-regulated in regenerative digits (amputation level < 33%) compared to non-regenerative digits (amputation level > 67%). Each sheet corresponds to a specific post-amputation time point.

**Supplementary Table S2:** Enriched GO terms for genes down-regulated in regenerative digits (amputation level < 33%) compared to non-regenerative digits (amputation level > 67%). Each sheet corresponds to a specific post-amputation time point.

### Supplementary Figures

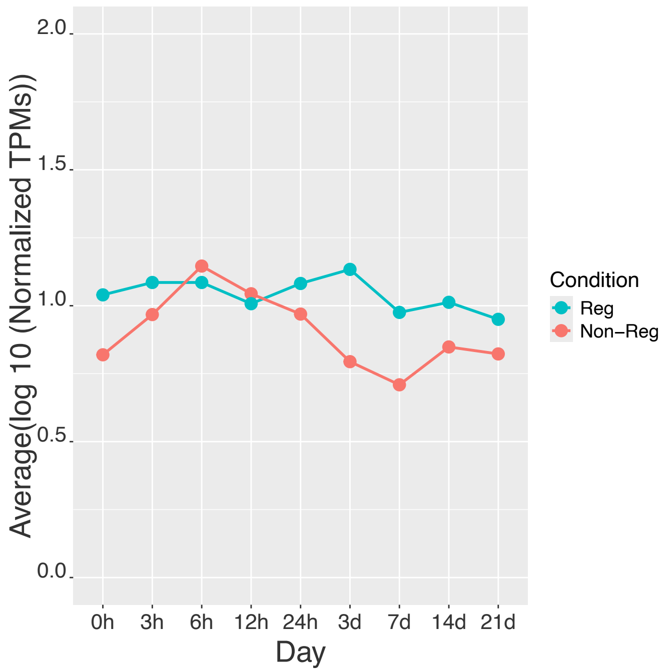


**Supplementary Figure S1:** Gene expression patterns of Bmp2 in regenerative and non-regenerative digits.
